# Pushing the limits of human motor control: high-frequency ankle oscillations in expert drummers

**DOI:** 10.1101/2025.06.18.660287

**Authors:** Aurélie Sarcher, Sylvain Dorel, Nadia R. Azar, Rémi Bal, Thomas Cattagni, Antoine Nordez

## Abstract

This study investigates the neuromechanical characteristics associated with drummers’ ability to achieve unilateral ankle oscillations with frequencies up to 10 Hz, surpassing known limits for lower-limb voluntary movements.

Eighteen experienced drummers performed rhythmic ankle oscillations across a range of imposed frequencies using a protocol combining motion capture and surface electromyography for trunk and lower limb muscles. Angular kinematics, muscle activation timing, and inter-muscle coordination were quantified to characterize neuromechanical adjustments associated with frequency modulation and to identify factors limiting maximal performance.

Drummers exhibited a preferred oscillation frequency of 6.3 Hz (range 5.5–7.3 Hz). At this frequency, soleus and tibialis anterior activation durations averaged 56.2 ms and 52.7 ms, respectively, and the latency from dorsiflexion to soleus activation was 48.5 ms. Increasing frequency was associated with shorter activation durations, reduced ankle excursion, and a shift toward plantar flexion. High-frequency performance involved substantial proximal muscle recruitment, particularly at the hip and trunk. Drummers achieving frequencies above 7.5 Hz (n = 6) displayed shorter tibialis anterior activation (34.3 vs. 53.2 ms) and lower tensor fascia latae activation (6.2% vs. 21.0%) compared with others.

High-frequency ankle oscillations in expert drummers exhibit neuromechanical features consistent with a constrained oscillatory system, reflecting interactions between limb mechanics, reflex pathways, and coordination constraints. Individual performance differences appear to depend on both distal muscle activation timing and proximal stabilization demands.

**Summary statement:** Expert drummers generate extremely rapid ankle oscillations through neuromechanical dynamics consistent with a constrained oscillatory system involving lower limb distal coordination and proximal stabilization.

## 1. Introduction

In metal music, expert drummers use a foot technique known as the ankle technique, which enables high-frequency bass drum strokes through oscillatory movements of a single leg. They commonly exceed 5 strokes per second unilaterally (i.e., 5 Hz) for minutes, with peaks of up to 10 Hz sustained briefly for the most skilled drummers. When combined with a double pedal setup (double bass drumming), the ankle technique is applied alternately with both legs, allowing drummers to double the number of bass drum strokes per second. While similar frequencies can be reached for the wrist and/or fingers (Aoki, Francis and Kinoshita, 2003; Aoki, Furuya and Kinoshita, 2005; Heuer, 2007; Fujii, Kudo, Ohtsuki, *et al*., 2009; Fujii, Kudo, Shinya, *et al*., 2009; Fujii and Moritani, 2012b, 2012a; Tanigawa *et al*., 2017; Eriksen *et al*., 2018; Beveridge *et al*., 2020; Sato *et al*., 2020), the frequencies typically observed lower limb movements during sports or music are significantly lower: even highly trained athletes rarely exceed 5 Hz, whether during full leg movements or ankle oscillations (Debraux and Bertucci, 2011; Tanigawa *et al*., 2017; Sato *et al*., 2020; Chaabouni *et al*., 2022).

Oscillations of the ankle with a frequency between 5 Hz and 10 Hz have been extensively documented, in the context of physiological tremor during voluntary movement (i.e., action or kinetic tremors) in healthy individuals (Stiles and Rietz, 1977; Iaizzo and Pozos, 1982; Pozos, Iaizzo and Petry, 1982; Pozos and Iaizzo, 1991; McAuley and Marsden, 2000; Deuschl *et al*., 2001; Jalaleddini *et al*., 2017), and of pathological tremor (i.e., clonus) observed in clinical populations (Pozos and Iaizzo, 1991; McAuley and Marsden, 2000; Deuschl *et al*., 2001; Beres-Jones, Johnson and Harkema, 2003). One well-supported explanation for dynamic ankle physiological tremor involves an interplay between mechanical resonance of the limb and spinal stretch reflex feedback, which may amplify oscillations under certain conditions (Raethjen *et al*., 2000; Deuschl *et al*., 2001, 2022). Consequently, the emergence and persistence of the oscillations are highly dependent on ankle angle, amplitude, and torque (Agarwal and Gottlieb, 1977; Stiles and Rietz, 1977; Pozos, Iaizzo and Petry, 1982; Evans *et al*., 1983; Rack *et al*., 1983; Jalaleddini *et al*., 2017). In addition, activation modulations of the soleus and tibialis anterior muscles, including onset timings and durations, fall within time ranges typically associated with stretch reflex responses (Agarwal and Gottlieb, 1977; Pozos, Iaizzo and Petry, 1982; Evans *et al*., 1983; Rack *et al*., 1983; Jalaleddini *et al*., 2017). While they share similar frequencies, it remains unknown whether the neuromechanical signatures of the ankle technique are consistent with those observed during tremor.

Although high-frequency performance is critical in metal music, expert drummers show considerable variability in their maximal achievable frequency. As frequency increases, ankle oscillations tend to shrink, and the pedal beater may no longer gain enough momentum to effectively strike the drum. Additionally, increasing movement frequency requires two primary neuromechanical adaptations, each associated with a risk factor for musculoskeletal disorders (da Costa and Vieira, 2010; Salvalaio *et al*., 2012; Azar, 2020, 2022; Tuček and Vaněček, 2020; Demissie, Bayih and Demmelash, 2024). The first comes from the repetitive and high-frequency nature of the ankle technique, suggesting very short activations from agonist and antagonist muscles controlling the ankle joint. When activation durations cannot be sufficiently reduced, excessive co-activation may occur, leading to excessive muscle tension, reduced mechanical efficiency and restricted movement range (Osu *et al*., 2002; Beveridge *et al*., 2020). Second, the need to stabilize the lower limb, pelvis and trunk segments during the ankle technique involves sustained isometric contractions of proximal muscles, particularly around the knee, hip and sacroiliac joints. These prolonged efforts can cause fatigue or pain due to elevated activation levels over time (Philipson *et al*., 1990; Levy *et al*., 1992; Bejjani, Kaye and Benham, 1996; Möller *et al*., 2018; Mizrahi, 2021). In response to these challenges, drummers may adopt distinct motor strategies to cope with both distal frequency and proximal stabilization demands. To our knowledge, no previous study analyzed these individual strategies, while they likely contribute to the observed differences in maximal performance frequency.

Using kinematic and electromyographic analyses, the first aim of this study was to characterize the neuromechanical signatures of high-frequency ankle oscillations in expert drummers and determine the extent to which these signatures overlap with mechanisms previously described in oscillatory ankle behaviors. The second aim of this study was to identify individual factors limiting frequency capacity in experienced drummers. We hypothesized that co-activation at the ankle and high proximal muscle activation would emerge at maximal frequency, contributing to reduced movement amplitude and discomfort. By characterizing these neuromechanical constraints, this study provides insights into how the human motor system manages high-frequency, high-intensity cyclic movements – informing strategies for injury prevention and performance optimization in drumming and other rhythmic motor tasks.

## 2. Materials and Methods

### 2.1 Participants

Eighteen male drummers were included in this study (age: 34 ± 8 years, height: 179 ± 8 cm, body mass: 76 ± 13 kg). Although inclusion was not restricted by sex, recruitment focused on drummers able to perform the ankle technique at an expert level and sustain very high-frequency double-bass drumming. Female drummers were actively sought during recruitment; however, no female drummer meeting the inclusion criteria could be identified within the recruitment period. This likely reflects the limited representation of women in extreme metal drumming and the relatively small proportion of drummers who master this highly specialized technique. Consequently, the findings should be interpreted as applying to the population studied and future research should investigate whether similar neuromechanical characteristics are observed in female drummers. Their anthropometric, drumming set-up, and drumming playing characteristics are summarized in Tables S1 and S2. All had mastered the ankle technique for at least 5 years. Participants’ drumming level varied between regional (performing frequently in shows, without participating in tours; n=7), national (performing frequently in shows and participating to tours of less than 15 consecutive shows; n=7), and international (performing frequently in shows and participating in international tours of more than 15 consecutive shows; n=4).

The study was approved by the local Research Ethics Boards of Nantes Université (reference number n°25112022) and the University of Windsor (reference number 23-077). Written informed consent was obtained for all participants, after they were informed about methods used in the study. All the experiments were conducted in accordance with the latest version of the Declaration of Helsinki and the Tri-Council Policy Statement: Ethical Conduct for Research Involving Humans (Canadian Institutes of Health Research, Natural Sciences and Engineering Research Council of Canada and Social Sciences and Humanities Research Council of Canada, 2022), which is based on the Code of Ethics of the World Medical Association (Declaration of Helsinki) for experiments involving humans.

### 2.2 Experimental protocol

The experimental protocol was divided into 2 parts: dynamic trials, and Maximal Voluntary Contractions (MVC) trials for proximal muscles.

#### 2.2.1 Dynamic trials

Figure 6 displays the drumming and experimental set-up for the dynamic trials.

##### Drumming set-up

The double bass drumming stand included a drum throne, two single pedals with two bass drum pads or double pedals with a single bass drum pad, a snare pad and a pair of sticks. Each pedal consists of a board connected to a beater, which strikes the bass drum when the board is pressed down by the foot. Once the pressure is released, a spring mechanism pulls the board and beater back to their initial positions, ready for the next strike. In the case of double pedals, a drive shaft connects the two pedals so that one pedal can control one of two beaters on the main pedal, allowing both beaters to strike the same bass drum.

Drummers came with their own equipment when possible, or adjusted all the settings of the equipment we provided to match their preferred settings. Those settings were collected for each drummer (i.e., drum throne: height; pedals: orientation of the beaters, tension of the springs, distance between the pedals, and board height. See Table S1.

##### Bass drum trigger

The signal from the bass drum electronic pad was collected at 2000 Hz by the motion analysis system.

##### Kinematics

Lower limb (bilateral) and trunk kinematics were recorded at a sampling frequency of 200 Hz using a 12-camera 3D motion analysis system (Vero 2.2 VICON, Oxford, UK) and software (Nexus 2.12.0, VICON, Oxford, UK). Thirty-six retro-reflective markers were placed on anatomical landmarks of the thorax, pelvis, thighs, legs, rear feet and forefeet, according to the two-segment foot pyCGM 2.4 model (Leboeuf, Baker, *et al*., 2019). Rear feet and forefeet markers, except for those on the lateral and medial malleoli, were placed directly on the drummers’ shoes, to preserve their playing habits.

##### Electromyography

Muscle activity was recorded at 2000 Hz using a wireless EMG system (PicoEMG, Cometa, Bareggio MI, Italy) and software (EMGandMotionTools 8.11.0.0, Cometa, Bareggio MI, Italy). EMG electrodes were equipped with Ag/AgCl electrodes with a recording diameter of 10 mm (Covidien, Mansfield, MA, USA), and placed bilaterally on seven muscles of the lower limbs and trunk: *soleus*, *tibialis anterior*, *rectus femoris*, *tensor fascia latae*, *gluteus medius*, *rectus abdominis*, and *erector spinae*. Pilot data using ultrasound measurements indicated that the gastrocnemius muscles contracted very little compared to the soleus muscle during the ankle technique, which is consistent with the knee flexed position of drummers. Therefore, the soleus and the tibialis anterior were considered as the main ankle plantar flexor and dorsiflexor muscles, respectively. Proximal muscles were selected based on discussions with drummers on principal pain locations: hip flexors, glutes, and trunk extensors stood out in most of the discussions. SENIAM electrode placement guidelines (Hermens *et al*., 1999) were followed for all muscles except for the soleus muscle, for which electrodes were positioned on the latero-posterior compartment based on ultrasound identification. The EMG system was triggered by the motion analysis system, ensuring a perfect synchronization of the kinematics and EMG data (<0.5 ms delay).

##### Procedure

First, drummers were asked to stand still to perform a static calibration of the kinematical model. Thereafter, drummers were asked to sit on their drum throne, and to warm up during a few minutes. To mimic the playing posture and ensure consistent trunk engagement, participants were asked to perform simple and repetitive arm movements on the snare pad during each trial. These movements consisted of basic rhythmic strokes, designed to simulate typical drumming posture without introducing complex upper-limb coordination or excessive variability. This ensured that trunk and upper-body muscle engagement remained representative of real drumming conditions, while keeping the primary focus on lower-limb neuromechanical analysis. Each drummer had their own range of frequencies for which they used the ankle technique, as well as a self-reported comfort frequency—i.e., the frequency at which they felt they could perform the longest with minimal muscle fatigue. In the drumming community, those frequencies are commonly presented in Beats Per Minute (BPM), with each beat corresponding to a quarter note. For clarity, all frequencies will be presented as unilateral frequencies in Hz. For example, a frequency of 180 BPM corresponds to a unilateral frequency of 6 Hz. Drummers were asked to perform trials of 20 seconds for several frequencies, with the tempo controlled by a metronome. After determining their own preferred frequency, the frequency of the metronome increased and then decreased by 0.33 Hz (equivalent to 10 BPM) up to the minimal and the maximal frequency they could reach. Each trial was spaced by two minutes of rest. Kinematic markers and EMG electrodes for tibialis anterior and soleus muscles were then removed.

#### 2.2.2 MVC trials

In this final phase of the protocol, drummers were placed on a clinical bed, still equipped with EMG electrodes on the proximal muscles. For EMG normalization purposes, they were asked to perform three isometric MVC, spaced by 2 minutes of rest, for each of the following conditions (Figure S1):

1. In a prone position, maximal trunk extensions were performed while being strapped to the bed at the shoulder level to normalize EMG signals from the left and right *erector spinae*.
2. In a supine position, maximal trunk flexions were performed while being strapped to the bed at the shoulder level to normalize EMG signals from the left and right *rectus abdominis*.
3. In a lateral position, maximal hip abductions were performed while being strapped to the bed at the knee level, to normalize EMG signals from the left and right *gluteus medius*.
4. In a seated position, maximal hip flexions were performed while being strapped to the bed at the thigh level to normalize EMG signals from the left and right *rectus femoris* and *tensor fascia latae*. Drummers could use their hands to stabilize their trunk, however they were not allowed to produce any force with the upper limbs.

### 2.3 Data processing

#### 2.3.1 Dynamic and MVC trials

Custom Matlab (Mathworks, Natick, MA, USA) routines written with the open-source Biomechanical Toolkit (Barre and Armand, 2014) were used to process the kinematic and EMG data.

##### Cycle detection

The signal from the bass drum trigger was used to detect the drumming cycles, which were defined as the time between two bass drum hits. Real trial frequency was then calculated using this data. Trials for which the real mean frequency deviated from the instructed frequency by more than 0.17 Hz (equivalent to 5 BPM) were considered as failed. The failed trial with the highest frequency was kept and defined as the maximal frequency trial for each participant. For each trial and each participant, timing parameters from EMG and kinematics were extracted from 12 cycles of movement.

##### Kinematics

The kinematics of the trunk, pelvis, hip, knee, and ankle joints were calculated using the pyCGM 2.4 model (Leboeuf, Baker, *et al*., 2019), available as an open-source Python package. The pyCGM 2.4 model is an evolution of the CGM model (Baker *et al*., 2017), and uses an inverse kinematic approach (Lu and O’Connor, 1999) with optimized hip joint center estimation (Hara *et al*., 2016; Leboeuf, Reay, *et al*., 2019). It also includes a two-segment foot model, with the rear segment adapted from the CGM, and the forefoot modified from the Oxford Foot Model (Carson *et al*., 2001).

##### Electromyography

Raw EMG signals were band-pass filtered (10-450 Hz, Butterworth zero-lag 4th order) and rectified. For the *tibialis anterior* and *soleus* EMG signals, the Teager-Kaiser energy operator was used to precisely detect the activation onset and offset timings (Solnik *et al*., 2008), using a semi-automatic threshold and a minimal window of 5 ms. For the proximal muscles, EMG signals were low-pass filtered (50 Hz, Butterworth zero-lag 2th order), and amplitude-normalized using the maximal activation measured in a 100 ms window of the MVC trials. 90^th^ percentile EMG amplitude levels were extracted from these normalized EMG signals, as typically done in ergonomic studies (Mathiassen, Burdorf and van der Beek, 2002; Graham, Agnew and Stevenson, 2009; Åkesson, Balogh and Hansson, 2012).

#### 2.3.2 Overview of the parameters collected

The following parameters were collected: duration between offset of tibialis anterior and onset of soleus (ms), duration between onset and offset of soleus (ms), duration between offset of soleus and onset of tibialis anterior (ms), duration between onset and offset of tibialis anterior (ms), duration between onset of ankle dorsiflexion and onset of soleus (ms), duration between onset of soleus and onset of ankle plantar flexion (ms), duration between onset of ankle plantar flexion and onset of tibialis anterior (ms), duration between onset of tibialis anterior and onset of ankle dorsiflexion (ms), ankle amplitude of plantar flexion / dorsiflexion (°), ankle mean position angle (°), 90^th^ percentile EMG amplitude levels of *tensor fascia latae*, *rectus femoris*, *gluteus medius*, *erector spinae* and *rectus abdominis* muscles (%). The durations between the onset of ankle dorsiflexion and soleus activation, as well as between the onset of ankle plantar flexion and tibialis anterior activation were extracted to characterize temporal relationships between movement and muscle activation, including time scales compatible with short-latency reflex responses (see Discussion 3.1). The durations between the onset of soleus activation and ankle plantar flexion, as well as between the onset of tibialis anterior activation and ankle dorsiflexion, were also calculated to describe the temporal coupling between muscle activation and joint movement, potentially reflecting muscle-tendon dynamics (see Discussion 3.1).

### 2.4 Data statistics

#### 2.4.1 Descriptive statistics

Mean and Standard Deviation (SD) of all variables were calculated for the 6.3 Hz trial, corresponding to the mean comfort frequency, for the 6.7 Hz trial, corresponding to the highest frequency common to all drummers, and for each drummer’s maximal frequency.

#### 2.4.2 Linear mixed-effects models: effect of the movement frequency

Linear mixed-effects models were used to analyze the effect of movement frequency on all parameters listed above (see 4.3.3), accounting for both fixed and random effects. The models were fitted using the lmer function from the lme4 package in R, with movement frequency as a fixed effect and random intercepts for subjects to account for repeated measures within individuals. Statistical significance of fixed effects was assessed using t-tests with Satterthwaite’s approximation for degrees of freedom, implemented via the lmerTest package. Model assumptions, including normality of residuals and homoscedasticity, were verified through diagnostic plots. The significance threshold was set at P<0.05.

#### 2.4.3 Comparison between drummers at common highest frequency

Participants were divided into two groups based on their maximal movement frequency: those above 7.5 Hz (n=6, group A) and those at or below 7.5 Hz (n=12, group B). Comparisons of 28 variables between the two groups, including 15 that were based on the highest common frequency of 6.7 Hz (Tables S1 to S5), were performed using the non-parametric Mann-Whitney U test, as the groups had unequal sizes. The significance threshold was set at P<0.05. The frequency threshold of 7.5 Hz was determined by testing multiple potential threshold values (from 7.0 to 7.9 Hz) and by assessing the stability of the P-values across these thresholds. The threshold for which the P-values remained significant across neighboring thresholds (above and below the selected threshold) was retained, ensuring the robustness of the results. Additionally, this choice was supported by the fact that, a frequency barrier, generally observed between 7.3 and 7.7 Hz in drummers, is difficult to exceed (Payne, no date), further reinforcing the relevance of the selected threshold.

## 3. Results

### 3.1 Dynamic trials

The following frequencies are unilateral movement frequencies, meaning that a drummer with a frequency of 5 Hz strikes the bass drum 10 times per second using both legs. Trial frequencies were calculated from the bass drum trigger and expressed unilaterally. Trials for all participants ranged from 4.6 Hz to 8.7 Hz. Participants did an average of 7 trials each (SD: 2), with a mean comfort movement frequency of 6.3 Hz (SD: 0.5 Hz), and a mean maximal frequency of 7.0 (SD: 1.8 Hz). The highest frequency common to all drummers was 6.7 Hz.

### 3.2 Objective # 1: Characterization of the ankle technique

#### 3.2.1 Timeline of the ankle technique and modulation of the parameters with movement frequency

To characterize the ankle technique, the results include ankle kinematics and physiological parameters extracted from the trial corresponding to the mean comfort movement frequency (i.e., 6.3 Hz), and slopes (fixed effects) of the linear mixed models, quantifying the effect of movement frequency on the different parameters. Figure 1 displays the fixed effect of the linear mixed-effects model analyzing effect of movement frequency on ankle mean position (°). Figures 2 and 3 display the ankle kinematics and physiological parameters results (respectively) with mean values for the trial at 6.3 Hz along with the effect of movement frequency on those parameters.

**Figure 1.**
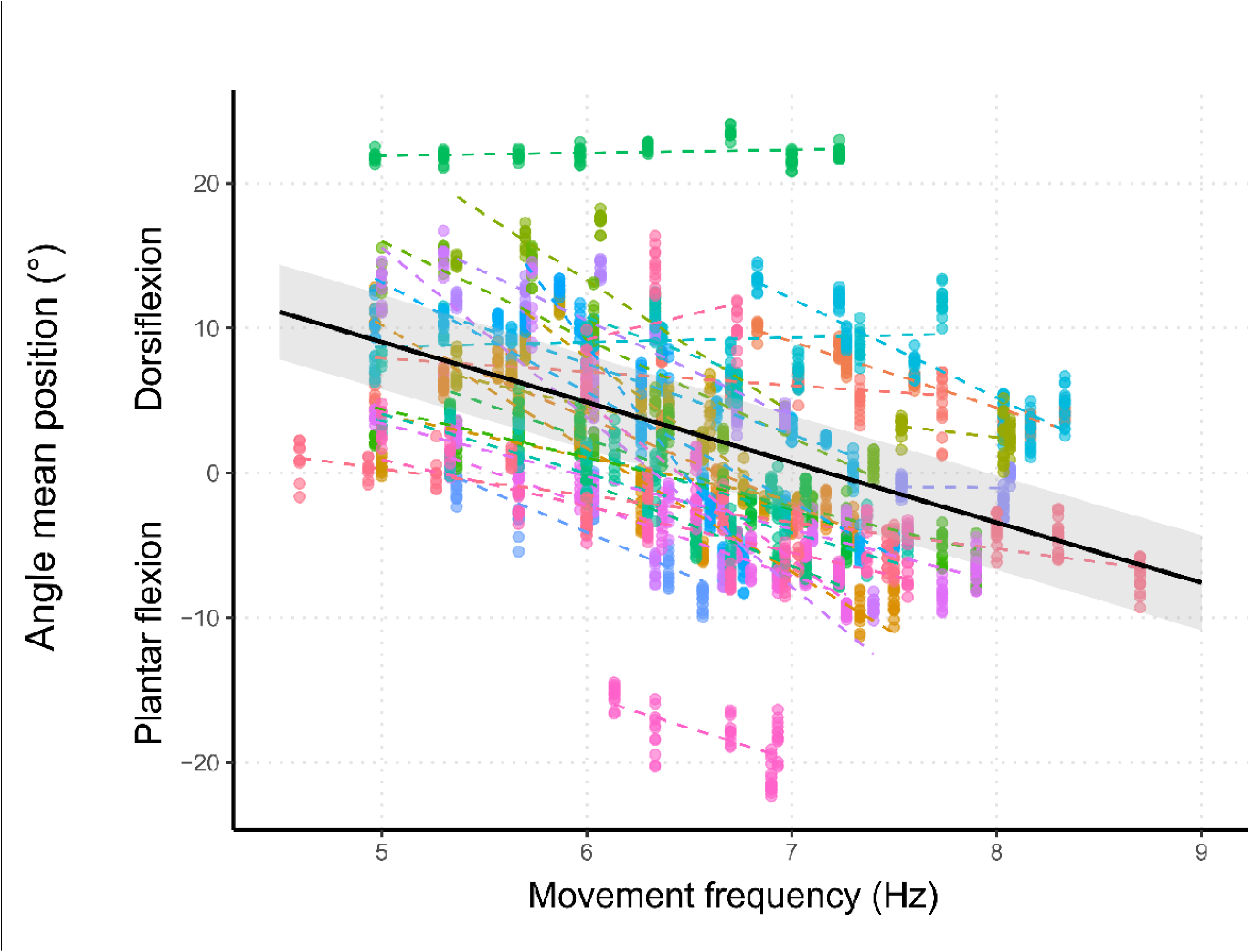
Fixed effect of the linear mixed-effects model on ankle position. The colored dots represent individual data points for each side of each subject, and the dashed lines show subject-specific regression trends. The solid black line represents the fixed effect of the linear mixed-effects model, with the shaded grey area indicating the 95% confidence interval. The slope of the black line corresponds to the fixed effect estimate from the model (slope = −4.2 °/Hz), reflecting the overall relationship between unilateral movement frequency and mean ankle position. A steeper slope would indicate a stronger association between frequency and ankle mean position across subjects. Positive values represent ankle dorsiflexion, negative values ankle plantarflexion, 0° represents the neutral position.

**Figure 2.**
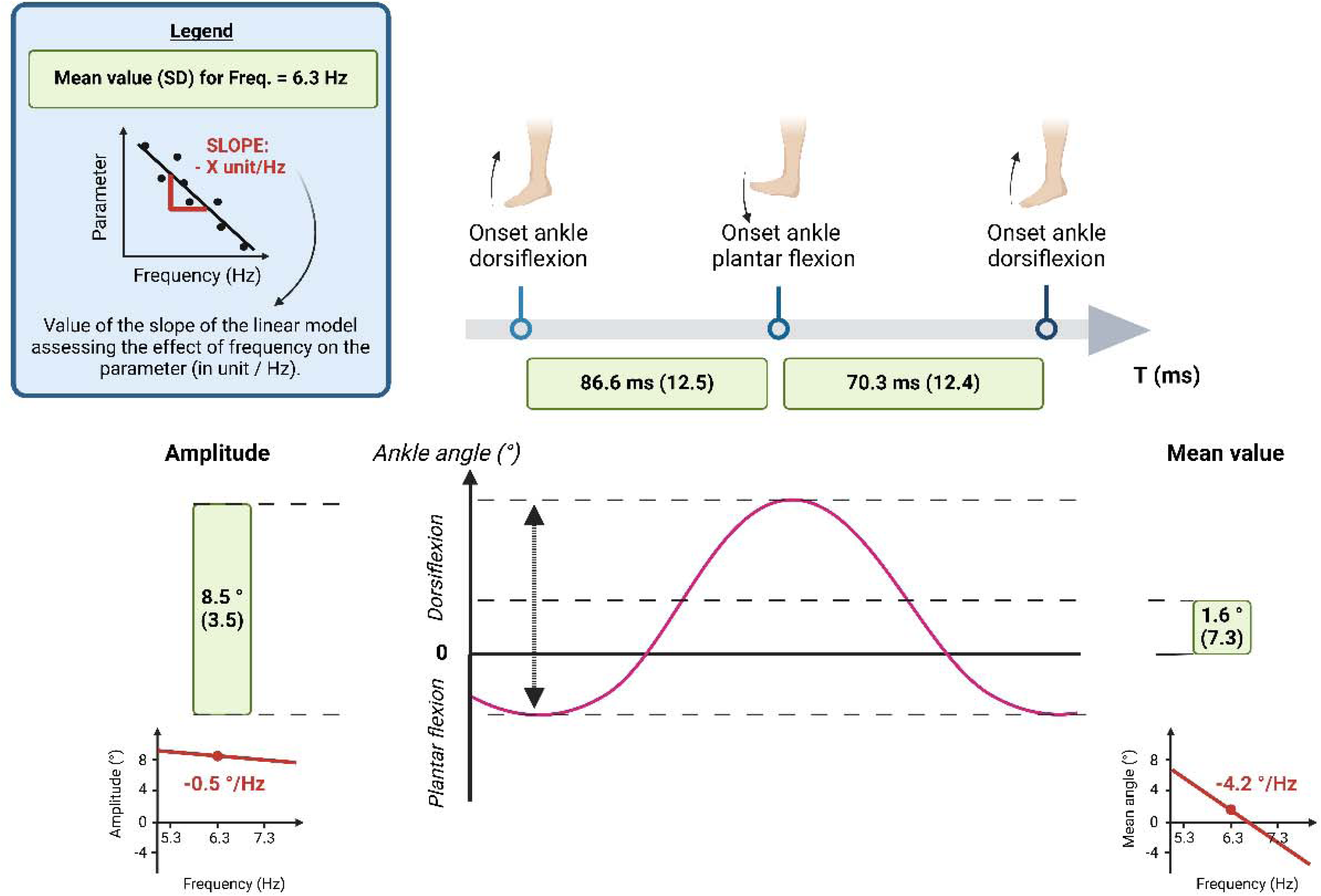
Kinematics parameters (amplitude and mean position of the ankle) during an ankle technique cycle. Framed values are absolute mean values (with standard deviation SD) for the trial at the frequency of 6.3 Hz. Linear mixed-effects models were used to analyze the effect of movement frequency on these parameters. Below each frame is indicated the slope of the movement frequency effect (unit/Hz).

**Figure 3.**
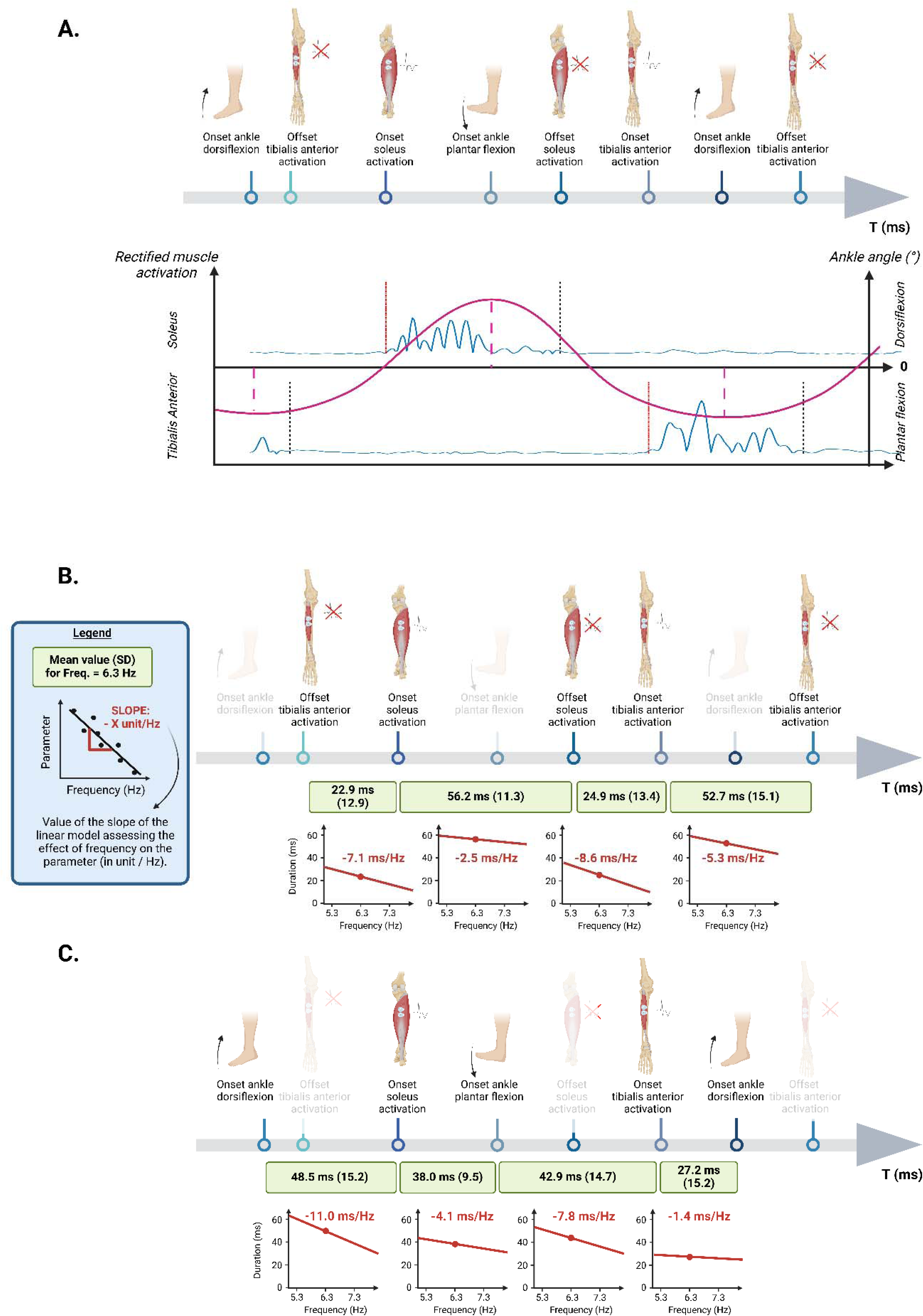
A. Rectified muscle activation signals are displayed for the soleus and tibialis anterior muscles, as well as ankle angle (°). The vertical lines correspond to onset (red, dash-dotted) and offset (black, dashed) detections using the Teager-Kaiser energy operator, and onset of ankle dorsiflexion / plantar flexion (magenta, large dashed). B. Timeline of the ankle technique, including analysis of muscle activation durations and off-periods for soleus and tibialis anterior muscles during an ankle technique cycle. C. Timeline of the ankle technique, including analysis of physiological parameters during an ankle technique cycle: durations between the onset of ankle dorsiflexion and soleus activation, as well as between the onset of ankle plantar flexion and tibialis anterior activation (to be compared to the short-latency stretch reflex), and durations between the onset of soleus activation and ankle plantar flexion, as well as between the onset of tibialis anterior activation and ankle dorsiflexion (used as a proxy of the electromechanical delay of the muscle-tendon unit). For B and C, framed values are absolute mean (with standard deviation SD) for the trial at 6.3 Hz. Linear mixed-effects models were used to analyze the effect of movement frequency on these parameters. Below each frame is indicated the slope of the movement frequency effect (unit/Hz). All models are statistically significant.

Regarding ankle kinematics (Figure 2), ankle amplitude of dorsiflexion / plantar flexion had a mean value of 8.5 ° (SD: 3.5°), and movement frequency had a small but significant negative impact on this value, with a slope of −0.5 °/Hz (F=108.7, dof=2728.7, P<0.001). Ankle mean position was 1.6 ° (SD: 7.3°), and movement frequency had a significant large negative impact on this value, with a slope of −4.2 °/Hz (F=1623.3, dof=2725.3, P<0.001). Thus, higher frequencies led to smaller amplitudes and more plantar flexion.

**Figure 3A shows the timeline of the ankle technique events.**

Regarding muscle activation (Figure 3B), duration of soleus activation had a mean value of 56.2 ms (SD: 11.3 ms), and movement frequency had a small but significant negative impact on this value, with a slope of −2.5 ms/Hz (F=81.0, dof=2737.1, P<0.001). Duration of tibialis anterior activation had a mean value of 52.7 ms (SD: 15.1 ms), and movement frequency had a significant moderate negative impact on this value, with a slope of −5.3 ms/Hz (F=230.8, dof=2254.6, P<0.001). Duration between offset of tibialis anterior and onset of soleus had a mean value of 22.9 ms (SD: 12.9 ms), and movement frequency had a significant large impact on this value, with a slope of −7.1 ms/Hz (F=405.3, dof=1984, P<0.001). Duration between offset of soleus and onset of tibialis anterior had a mean value of 24.9 ms (SD: 13.4 ms), and movement frequency had a significant large negative impact on this value, with a slope of −8.6 ms/Hz (F=568.2, dof=2251, P<0.001). These findings indicate that with increasing frequency, activation durations shortened, and off-periods reduced even more.

Regarding the temporal relationships between joint movement and muscle activation (Figure 3C), duration between the onset of ankle dorsiflexion and soleus activation had a mean value of 48.5 ms (SD: 15.2 ms), and movement frequency had a significant large negative impact on this value, with a slope of −11 ms/Hz (F=1108.2, dof=2735.7, P<0.001). Duration between the onset of ankle plantar flexion and tibialis anterior activation had a mean value of 42.9 ms (SD: 14.7 ms), and movement frequency had a significant large negative impact on this value, with a slope of −7.8 ms/Hz (F=564.3, dof=2256, P<0.001).

Regarding the temporal delays between muscle activation and joint movement (Figure 3C), duration between the onset of soleus activation and ankle plantar flexion had a mean value of 38.0 ms (SD: 9.5 ms), and movement frequency had a significant moderate negative impact on this value, with a slope of −4.1 ms/Hz (F=388, dof=2735.7, P<0.001). Duration between the onset of tibialis anterior activation and ankle dorsiflexion had a mean value of 27.2 ms (SD: 15.2 ms), and movement frequency had a small but significant negative impact on this value, with a slope of −1.4 ms/Hz (F=12.8, dof=2253.9, P<0.001).

As movement frequency increased, drummers were able to decrease all measured physiological parameters. However, the durations between the onset of soleus and tibialis anterior fiber stretching and their respective activation were reduced more than the proxies of the electromechanical delay.

### 3.3 Objective # 2: Individual characterization of drummers

Participating drummers’ individual characteristics and results are displayed in Tables S1 to S5, for each drummer’s maximal movement frequency, and for the highest common frequency of 6.7 Hz.

#### 3.3.1 Individual analysis at maximal frequency

At maximal frequency, 6 out of 18 drummers displayed durations between offset of tibialis anterior or soleus and onset of soleus or tibialis anterior that were close to, or lower than, zero (Table S3), indicating temporal overlap between soleus and tibialis anterior activations, and therefore co-activation. This phenomenon can be observed in Figure 4, which displays the soleus and tibialis anterior activations for participants 13 and 11: at their maximal frequency (respectively 7.2 Hz and 7.9 Hz), duration between offset of soleus and onset of tibialis anterior is only 5 ms for participant 13 (Figure 4D), whereas it is 24 ms for participant 13 (Figure 4A).

**Figure 4.**
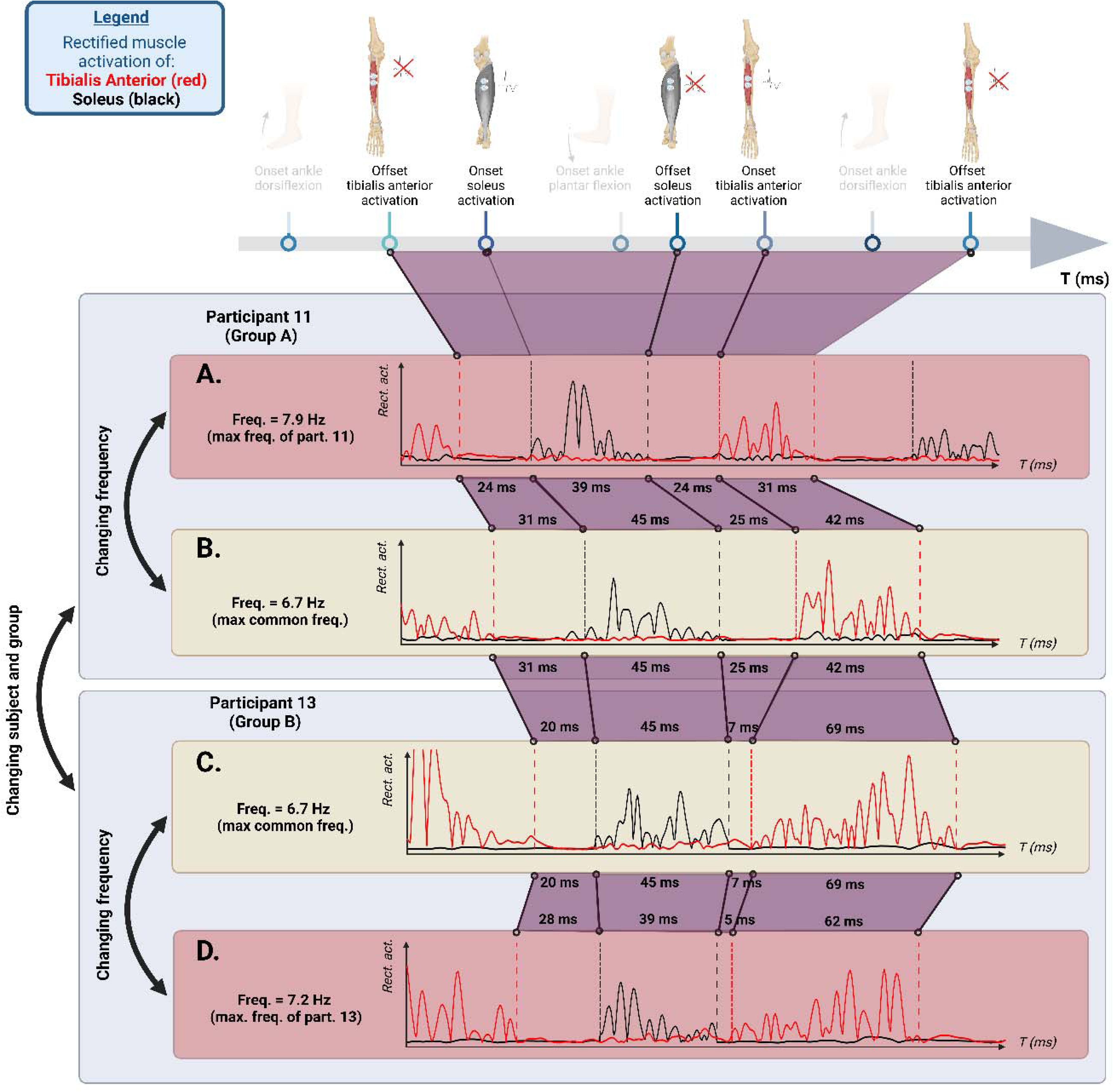
Rectified muscle activation of right soleus (black) and tibialis anterior (red) muscles for A. participant 11 from group A (maximal frequency > 7.5 Hz) at his maximal frequency of 7.9 Hz, B. participant 11 from group A at the highest common frequency of 6.7 Hz, C. participant 13 from group B (maximal frequency ≤ 7.5 Hz) at the highest common frequency of 6.7 Hz, D. participant 13 from group B at his maximal frequency of 7.2 Hz. Values (ms) are indicated between offset of tibialis anterior and onset of soleus, between offset of soleus and onset of tibialis anterior, and for the activation duration of soleus and tibialis anterior.

Additionally, at maximal frequency, 13 out of 18 drummers had values of the 90^th^ percentile activation of at least one proximal muscle higher than 30% MVC (Maximal Voluntary Contraction) (Table S5), indicating high levels of activation in at least one of the muscles stabilizing the trunk, pelvis, or upper leg segments. This phenomenon can be observed in Figure 5, which displays the mean and standard deviation values for the 90^th^ percentile activation of the five measured proximal muscles for participants 13 and 11: at their maximal frequency (respectively 7.2 Hz and 7.9 Hz), participant 13 displays high levels of activation in tensor fascia latae, rectus femoris, erector spinae and gluteus medius (Figure 5D), and participant 11 displays high levels of activation in rectus femoris and gluteus medius (Figure 5B).

**Figure 5.**
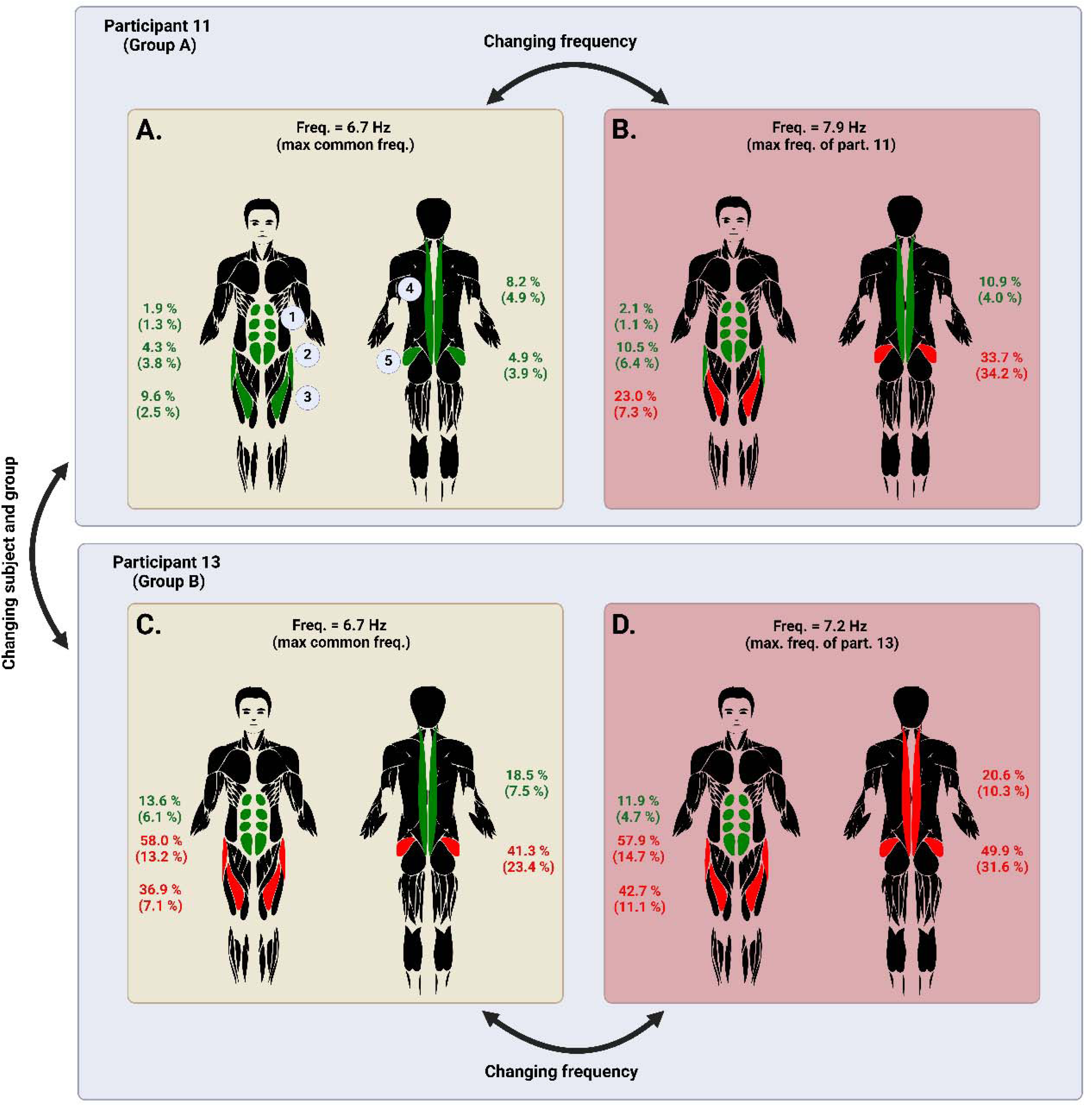
Mean (Standard Deviation) values for the 90^th^ percentile of muscle activation in percent of the Maximal Voluntary Contraction (MVC), averaged for both sides, for rectus abdominis (1), tensor fascia latae (2), rectus femoris (3), erector spinae (4) and gluteus medius (5) muscles, for A. participant 11 from group A (maximal frequency > 7.5 Hz) at the highest common frequency of 6.7 Hz, B. participant 11 from group A at his maximal frequency of 7.9 Hz, C. participant 13 from group B (maximal frequency ≤ 7.5 Hz) at the highest common frequency of 6.7 Hz, D. participant 13 from group B at his maximal frequency of 7.2 Hz. Values (%) and localizations of each studied muscle are indicated in green if mean+SD < 30 % MVC, and in red if mean+SD > 30 % MVC.

**Figure 6.**
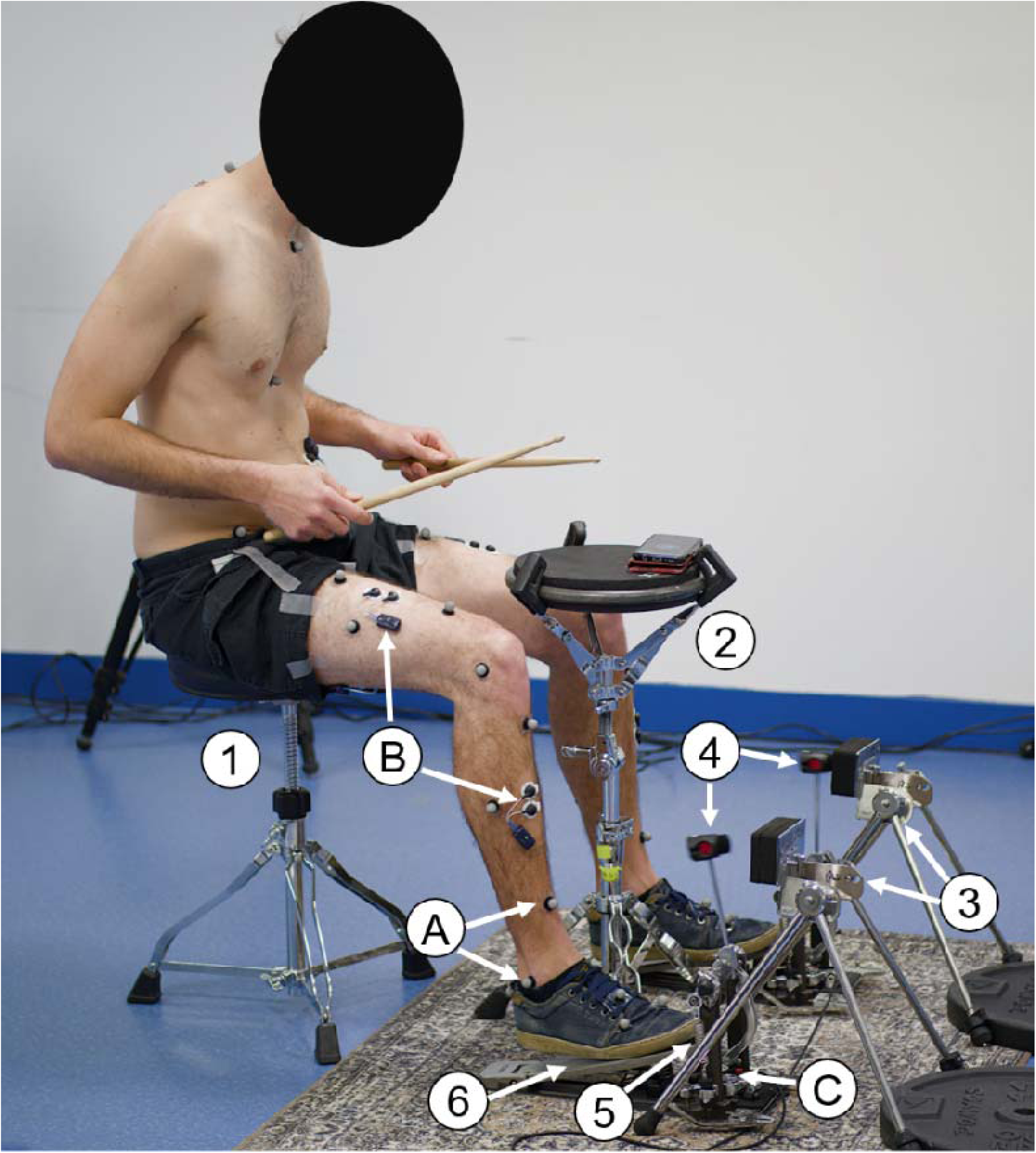
Drumming and experimental set-up for the dynamic trials: 1. Drum throne, 2. Snare pad, 3. Bass drum pad, 4. Pedal’s beater, 5. Pedal’s spring, 6. Pedal’s board, A. Motion capture markers, B. Electromyography electrodes, C. Bass drum trigger.

Those two phenomena of co-activation between soleus and tibialis anterior activations, and high levels of activation in proximal muscles could be limiting factors for the drummer’s maximal frequency.

#### 3.3.2 Comparison between drummers at highest common frequency

Participants with a maximal frequency higher than 7.5 Hz (Group A; n=6; drummers [1; 2; 8; 11; 16; 18]; mean maximal frequency=8.1 Hz, SD: 0.4 Hz) displayed several differences with the other participants (Group B; n=12; mean maximal frequency=7.1 Hz, SD: 0.3 Hz), for the trial at the highest common frequency of 6.7 Hz. These differences may explain why some drummers reach higher frequencies.

Group A had shorter duration of activation of the tibialis anterior (mean=34.3 ms, SD: 5.8 ms) than group B (mean=53.2 ms, SD:14.1 ms) (U=17; P=0.0013), shorter duration between onset of tibialis anterior and ankle dorsiflexion (mean=17.5 ms, SD: 6.0 ms) than group B (mean=28.7 ms, SD: 10.2 ms) (U=25; P=0.0365), and lower level of tensor fascia latae activation (mean=6.2 %, SD: 2.8%) than group B (mean=21.0 %, SD: 16.3%) (U=31; P=0.0135). Duration between offset of soleus and onset of tibialis anterior was slightly higher in group A (mean=32.4 ms, SD:7.4 ms) than group B (mean=23.1 ms, SD:11.0 ms), however this result was not statistically significant (U=60; P=0.1296). No differences were shown for all other parameters, including duration between onset of soleus and onset of ankle plantar flexion (U=47; P=0.3714), duration between offset of tibialis anterior and onset of soleus (U=55; P=0.3161), and spring tension (U=63; P=0.5956). These results indicate that drummers with the highest maximal frequency can produce shorter tibialis anterior activation, resulting in a larger temporal window before experiencing co-activation. Moreover, their timing of tibialis anterior activation is adapted, suggesting and adaptation of activation timing relative to the mechanical behavior of the pedal spring, but without any major spring tension differences. The differences in muscle activation durations highlighted at the highest common frequency of 6.7 Hz between participants from groups A and B can be observed in Figure 4: participant 11 from group A displays, in comparison to participant 13 from group B, shorter activation of tibialis anterior (42 ms versus 69 ms), longer duration between offset of soleus and onset of tibialis anterior (25 ms versus 7 ms), and longer duration between offset of tibialis anterior and onset of soleus (31 ms versus 20 ms) (Figure 4B and 4C), however this last individual difference was not highlighted by statistical testing at the groups level. The differences in tensor fascia latae levels highlighted at the highest common frequency of 6.7 Hz between participants from groups A and B can be observed in Figure 5: participant 11 from group A displays a mean 90^th^ percentile of 4.3 %, while participant 13 from group B displays a mean 90^th^ percentile of 58.0 % (Figure 5A and 5C). Additionally, in comparison to participant 13 from group B, participant 11 from group A displays a much lower 90^th^ percentile activation of rectus femoris (9.6 % versus 36.9 %) and gluteus medius (4.9 % versus 41.3 %) (Figure 5A and 5C), however these last individual differences were not highlighted by statistical testing at the groups level.

## 4. Discussion

The results of the present study support our first hypothesis by showing that the ankle technique used by expert drummers exhibits neuromechanical features consistent with a constrained oscillatory system operating at the ankle joint. In particular, movement frequency, ankle posture, oscillation amplitude, and muscle activation timing were modulated in ways that overlap with patterns reported in the literature on ankle oscillations observed during tremor. In accordance with our second hypothesis, we also observed marked interindividual variability in distal and proximal muscle activation patterns, which appears to be associated with the ability to perform the ankle technique at higher frequencies. Taken together, these findings suggest that high-frequency bass drum strokes are not achieved solely through cycle-by-cycle voluntary control, but emerge from the interaction between neural control, lower limb–pedal closed loop system, and individual motor strategies. This perspective helps explain both performance variability and the musculoskeletal demands associated with the technique.

### 4.1 The ankle technique as a constrained neuromechanical oscillator

First, our results showed that drummers had a mean comfort movement frequency of ankle oscillations of 6.3 Hz, in the range of commonly frequencies reported for tremor, where typical values lie between 4 and 8 Hz (Agarwal and Gottlieb, 1977; Stiles and Rietz, 1977; Pozos, Iaizzo and Petry, 1982; Evans *et al*., 1983; Rack *et al*., 1983; Pozos and Iaizzo, 1991). The existence of an optimal movement frequency with a “natural resonance” of the ankle, associated to a specific plantar flexed ankle position, was also reported in previous ankle oscillations studies (Stiles and Rietz, 1977; Pozos, Iaizzo and Petry, 1982; Evans *et al*., 1983; Jalaleddini *et al*., 2017). Drummers were able to voluntarily modulate the oscillations frequency by 2 Hz, same as physiological oscillations (Stiles and Rietz, 1977; Deuschl *et al*., 2022). This modulation to higher frequencies was associated with reduced amplitude and a more plantar-flexed mean position, consistent with the literature (Stiles and Rietz, 1977), which also reports modulation of the torque applied to the ankle (Stiles and Rietz, 1977; Pozos, Iaizzo and Petry, 1982; Evans *et al*., 1983; Rack *et al*., 1983). This suggests that ankle configuration promotes different combinations of muscle-tendon unit mechanics and neural reflex factors, that change the resonance frequency of the limb, and therefore the oscillations frequency (Agarwal and Gottlieb, 1977; Stiles and Rietz, 1977; Pozos, Iaizzo and Petry, 1982; Evans *et al*., 1983; Jalaleddini *et al*., 2017).

Regarding mechanical factors, previous work has shown that oscillatory ankle behavior is strongly influenced by joint configuration and loading conditions, including ankle position, oscillation amplitude, and torque production (Agarwal and Gottlieb, 1977; Stiles and Rietz, 1977; Pozos, Iaizzo and Petry, 1982; Evans *et al*., 1983; Rack *et al*., 1983; Jalaleddini *et al*., 2017). Although our study did not measure muscle-tendon stiffness directly, it included the measurement of the delay between the onset of soleus activation and ankle plantar flexion (38.0 ms at the mean comfort frequency of 6.3 Hz), which is partly related to muscle-tendon unit length and stiffness (Kubo *et al*., 2001; Grosset *et al*., 2009; Longo *et al*., 2017). This delay significantly decreased with movement frequency, together with the shift toward a more plantar-flexed ankle position and the reduction in oscillation amplitude. In addition, previous studies have shown that ankle torque increases with oscillation frequency (Stiles and Rietz, 1977; Pozos, Iaizzo and Petry, 1982; Evans *et al*., 1983; Rack *et al*., 1983). Taken together, these findings suggest that increasing movement frequency modifies the mechanical operating conditions of the ankle-pedal system, leading to tighter temporal coupling between muscle activation and joint motion. Overall, these results are consistent with the increased mechanical constraints imposed by rapid oscillatory behavior.

Because detailed measurements of muscle activation during cyclic ankle oscillations remain limited in physiological conditions, some comparisons rely on studies conducted in pathological contexts, such as stretch-induced clonus, where resulting ankle oscillations share overlapping mechanical and physiological characteristics (Stiles and Rietz, 1977; Pozos, Iaizzo and Petry, 1982; Pozos and Iaizzo, 1991; Deuschl *et al*., 2022), despite different origins. In this context, soleus mean duration of activation was 56.2 ms at the mean comfort frequency of 6.3 Hz, similar to the value of 58 ms found by Beres-Jones et al. (Beres-Jones, Johnson and Harkema, 2003) during stretch induced clonus, at a mean frequency of 5.3 Hz. Tibialis anterior mean duration of activation was 52.7 ms at the mean comfort frequency of 6.3 Hz, slightly higher than the value of 43 ms found by Beres-Jones et al. (Beres-Jones, Johnson and Harkema, 2003) during stretch induced clonus, yet for a higher frequency of 7.3 Hz. Both soleus and tibialis anterior activation durations decreased significantly as movement frequency increased, as found in the tremor literature (Agarwal and Gottlieb, 1977; Stiles and Rietz, 1977; Pozos, Iaizzo and Petry, 1982; Evans *et al*., 1983; Rack *et al*., 1983; Pozos and Iaizzo, 1991), as well as in studies of high-frequency wrist movements in expert drummers (Fujii, Kudo, Ohtsuki, *et al*., 2009; Fujii, Kudo, Shinya, *et al*., 2009; Fujii and Moritani, 2012a; Beveridge *et al*., 2020), suggesting that compression of activation timing is a general requirement for rapid rhythmic performance. In our study, increasing movement frequency progressively compressed the temporal organization of the activation cycle. Although soleus and tibialis anterior activation durations became shorter, the silent intervals separating the two muscles decreased even more markedly, eventually leading to temporal overlap and co-activation in 6 of the 18 drummers at maximal frequency. Similar behavior was found in drummers during high-frequency wrist movements, with increased levels of co-activation with movement frequency (Fujii, Kudo, Ohtsuki, *et al*., 2009; Fujii, Kudo, Shinya, *et al*., 2009; Fujii and Moritani, 2012a; Beveridge *et al*., 2020). This indicates that the transition to higher frequencies may be limited not only by the ability to shorten agonist and antagonist bursts, but also by the ability to preserve sufficient temporal separation between them.

The delay between dorsiflexion onset and soleus activation averaged 48.5 ms at the mean comfort frequency of 6.3 Hz, which falls within the time range typically associated with short-latency stretch-related feedback at the ankle, between 40 and 50 ms (Agarwal and Gottlieb, 1977; Blackburn, Padua and Guskiewicz, 2008; Cronin *et al*., 2008). This observation indicates that the time scale of the movement is compatible with reflex loop dynamics. The fact that this delay decreased significantly with movement frequency is also noteworthy, and may reflect frequency-dependent changes in ankle configuration, mechanical coupling, or afferent sensitivity. Previous studies suggest that physiological tremor may vary with muscle shortening in ways that depend on afferent regulation rather than simple mechanical stretch alone (Stiles and Rietz, 1977; Pozos, Iaizzo and Petry, 1982; Jalaleddini *et al*., 2017): as movement frequency increases, the shift in ankle position towards greater plantar flexion enhances gamma fusimotor activity, which in turn increases muscle spindle sensitivity and amplifies the muscle’s response to stretching. Although presynaptic inhibition usually reduces the short-latency stretch reflex during voluntary movement, several studies show that this inhibition can decrease when strong cyclic Ia input is present, as in physiological tremor and clonus (Stiles and Rietz, 1977; Dimitrijevic, Nathan and Sherwood, 1980; Pozos and Iaizzo, 1991; Beres-Jones, Johnson and Harkema, 2003; Deuschl *et al*., 2022). Such conditions may transiently enhance reflex gain. Our results are consistent with the idea that stretch-related feedback may contribute to the oscillatory state, while not allowing its contribution to be isolated from mechanical or central factors.

Two additional classes of mechanisms should also be considered (McAuley and Marsden, 2000; Deuschl *et al*., 2001, 2022). First, central pattern generators can produce rhythmic output without sensory input (Grillner, Perret and Zangger, 1976; Grillner, 1985; Grillner and Wallén, 1985). However they generally offer limited flexibility in frequency modulation (Dimitrijevic, Nathan and Sherwood, 1980; Beres-Jones, Johnson and Harkema, 2003; Deuschl *et al*., 2022), which does not fit well with the voluntary ∼ 2 Hz modulation observed here and in physiological action tremor studies. Second, coupled oscillator models have been used to account for coordination between limb segments or muscle groups, particularly in tremor and locomotor contexts (Kelso, 1984; Haken, Kelso and Bunz, 1985; Hurtado *et al*., 2000; Huang and Ferris, 2010). In our case, however, the ankle technique can also be performed unilaterally, suggesting that the essential oscillatory behavior may arise within a single limb interacting with the pedal system. Bilateral drumming in our study reproduced real performance demands experienced by drummers, including trunk stabilization, offering an ecologically valid framework to assess injury risks and neuromechanical constraints.

Overall, this study shows that the ankle technique displays neuromechanical signatures consistent with a constrained oscillatory system, in which ankle mechanics, muscle-tendon behaviour, reflex-related feedback, and voluntary control may all interact to support high-frequency performance.

### 4.2 Individual neuromechanical factors associated with maximal frequency

Our individual analysis revealed that several mechanical and physiological parameters may influence maximal frequency.

Firstly, drummers that reached higher maximal frequencies had shorter and later activations of their tibialis anterior at the maximal common frequency of 6.7 Hz. This suggests that tibialis anterior timing plays an important role in regulating the return phase of the beater and in preserving a sufficient temporal window before co-activation with soleus occurs. We also observed a non-significant tendency for a longer interval between soleus offset and tibialis anterior onset in the higher-frequency group, which is directionally consistent with this interpretation. Together, these findings suggest that drummers who reach the highest frequencies may do so by compressing tibialis anterior activity while preserving the timing relationships needed to avoid excessive overlap between antagonist bursts. Similar associations between co-activation and performance have been reported in high-frequency wrist movements (Fujii, Kudo, Ohtsuki, *et al*., 2009; Fujii, Kudo, Shinya, *et al*., 2009; Fujii and Moritani, 2012a; Beveridge *et al*., 2020), where lower co-activation is linked to higher maximal frequencies, suggesting that minimizing antagonist overlap may be a general strategy for rapid rhythmic tasks.

Second, drummers who reached higher maximal frequencies displayed lower EMG activity levels for proximal muscles as the tensor fascia latae (6.2% versus 21%) at the maximal common frequency, indicating an ability to perform the ankle technique in a more relaxed state in proximal regions. Similar results were found for expert pianists that were more relaxed in proximal muscles than beginners (Furuya and Kinoshita, 2008). The ankle technique seems to carry increased risk for development of musculoskeletal disorders, especially at maximal frequency, since 13 of the 18 drummers displayed muscle activations higher than 30% MVC for at least one proximal muscle among the erector spinae, the gluteus medius, the rectus femoris and the tensor fascia latae. Such high activation levels reflect the need for proximal stabilization, as minimal leg weight is applied to the pedals. In line with this, it has been highlighted in the literature that drummers are strongly affected by playing-related musculoskeletal disorders, mostly in the upper limb and lower back regions (Roseiro *et al*., 2018; Azar, 2020, 2022). Our study therefore highlights the specificity of ankle technique in the development of musculoskeletal disorders in the lower back and hip regions.

Third, additional individual factors may affect maximal frequency even though they were not directly demonstrated here. Because oscillatory ankle behavior is influenced by muscle-tendon mechanics, it is plausible that tendon stiffness, passive ankle properties and pedal mechanics could affect the frequency range accessible to each drummer. In our study, we did not observe group differences in the delay between soleus activation and plantar flexion, suggesting that this temporal measure alone does not account for performance differences. However, we expect that a shorter or stiffer tendon could result in a shorter electromechanical delay (Kubo *et al*., 2001; Grosset *et al*., 2009; Longo *et al*., 2017), increased firing of muscle spindles (Abbott *et al*., 2024), therefore higher maximal frequency (Agarwal and Gottlieb, 1977). Likewise, we found no association between spring tension and maximal frequency, although the mechanics of the pedal system likely remain an important component of the overall oscillator, in relation with the lower limb inertia of the drummer (Evans *et al*., 1983): since the spring tension determines the force required to activate the pedal, as well as the return force and speed of the beater to its initial position, its setting could be related to the lower limb mass of the drummer. Because the drummer-throne-pedal system forms a closed loop, parameters such as pedal tension, limb inertia, and posture probably interact, and may need to be considered together rather than in isolation.

Finally, because the oscillatory regime appears to depend on both distal timing and proximal stabilization, training strategies may need to address both components. While strengthening exercises could improve proximal stabilization, it is possible that training affecting soleus muscle-tendon properties (Kubo, Ikebukuro and Yata, 2021; Ramírez-delaCruz *et al*., 2022) and reflex-related responsiveness (Voigt, Chelli and Frigo, 1998; Piirainen *et al*., 2014), such as plyometric training, may influence the ease with which drummers access this high-frequency oscillatory regime, although this remains to be tested directly.

### 4.3 Limitations of the study

Several limitations should be acknowledged. First, the temporal measures analyzed in this study were short and their resolution was relative to acquisition frequency, ranging from 0.5 ms for physiological events (electromyography at 2000 Hz) to 5 ms for kinematical events (motion capture at 200 Hz). Although our values fall within ranges reported in the literature, they should still be interpreted with appropriate caution.

Second, the results presented here are averaged for the left and right sides, whereas asymmetry during ankle technique can be relatively common in drummers, who often describe a “leading” and a “following” side. This likely contributed to the variability observed in several parameters. Because the present study focused on global characterization of the technique, such asymmetries were not analyzed separately, but they represent an important direction for future work.

Third, although the study identified timing relationships compatible with oscillatory ankle behavior and reflex-related time scales, it does not allow the respective contributions of mechanical resonance, feedback pathways, and central control processed to be disentangled. The discussion of these mechanisms should therefore be understood as an interpretation framework rather than a direct demonstration.

Finally, only a subset of proximal muscles was recorded. While musculoskeletal disorders in drummers have been referenced (Roseiro *et al*., 2018; Azar, 2020, 2022), to date, no literature specifically addresses the ankle technique. Muscle selection was guided by drummers’ reports of pain locations, which most often involved hip flexors, gluteal muscles, and trunk extensors. Other muscles may also contribute to proximal stabilization during the ankle technique. Some drummers also reported symptoms suggestive of piriformis syndrome, but its activation could not be measured using surface EMG.

## Conclusion

This study provides new insights into the biomechanics and neuromuscular control of the ankle technique used by drummers to achieve high-frequency bass drum strokes.

Our findings show that this technique exhibits neuromechanical features consisted with a constrained oscillatory system, including frequency-dependent changes in ankle posture, reduced movement amplitude at higher speeds, and compressed distal activation timing. These features overlap with patterns described in the literature on oscillatory ankle behavior, including physiological tremor and clonus, but do not imply a single underlying mechanism.

The ability to reach very high maximal frequencies appears to depend on individual motor strategies, particularly shorter and better-timed tibialis anterior activation and lower proximal muscle activation, especially of the tensor fascia latae. Thus, successful performance seems to rely on a delicate balance between distal timing precision, avoidance of excessive co-activation, and economical proximal stabilization.

Beyond performance, the ankle technique may also carry specific musculoskeletal demands, particularly in the lower back and hip regions, because of the high stabilization requirements imposed by the task. Future studies should investigate training adaptations, side asymmetries, and the interactions between distal oscillatory control and proximal stabilization to better understand both performance optimization and injury prevention.

## Supporting information

Supplementary Video 1 Ankle Technique

Supplementary Video 2 Swivel Technique

## Author Contributions

A.S. and A.N. conceived the experiment. A.S. and R.B. carried out the experiments and processed the data. A.S. interpreted the data in consultation with S.D., T.C. and A.N, and N.R.A.. A.S. wrote the paper with input from all authors who provided critical feedback and helped shape the manuscript.

## Acknowledgements

None.

## Data availability

The datasets generated and analysed during the current study are publicly available in the French Recherche Data Gouv repository at: https://doi.org/10.57745/THHWP2

## Consent to participate

Written informed consent was obtained from all participants.

## AI use

AI-based language assistance tools (ChatGPT OpenAI, GPT-5) were used for English editing and manuscript preparation. All scientific content, interpretations, and final text validation were performed by the authors.

## Abbreviations

EMG: electromyography
MVC: maximal voluntary contraction
SOL: soleus
TA: tibialis anterior
BPM: beats per minute

