## Supporting Information for "Pushing the limits of human motor control: high-frequency ankle oscillations in expert drummers"

**This PDF file includes:**

Figures S1
Tables S1 to S5
SI References

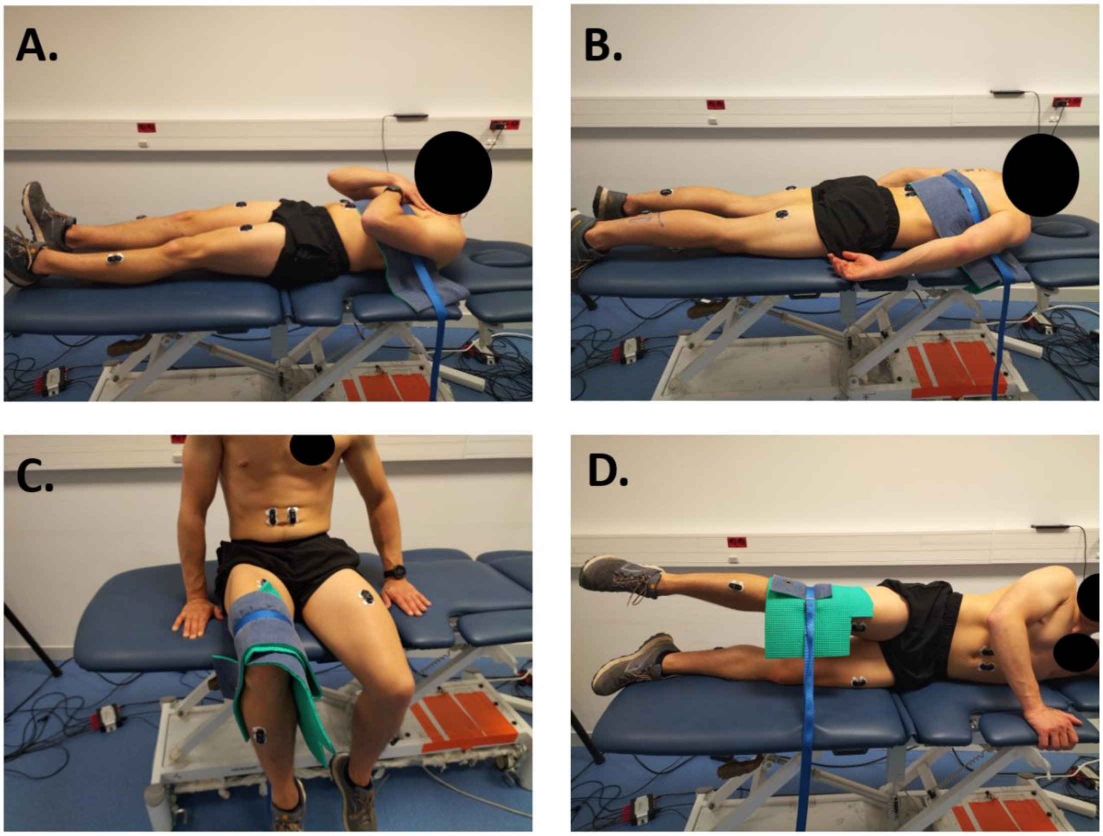


**Fig. S1.** Maximal Voluntary Contractions (MVC) trials, for A. trunk flexion, B. trunk extension, C. hip flexion, D. hip abduction.

**Table S1.** Participating drummers' anthropometric and drumming set-up characteristics. 1: Leg length was measured standing up, with a direct line from anterior superior iliac spine to medial malleolus. 2: Leg mass is 16.68% of the total body mass (1). 3: Throne height is the averaged height between the top of the cushion and the bottom of the cushion. Drummers were grouped according to their maximal frequency (i.e., A: > 7.5 Hz; B: ≤ 7.5Hz).

| | Anthropometric characteristics | | | | | Drumming set-up characteristics | | | | | |
|---|---|---|---|---|---|---|---|---|---|---|---|
| **Drummer – group** | **Age (years)** | **Height (cm)** | **Body mass (kg)** | **Leg length [1] (cm)** | **Leg mass [2] (kg)** | **Throne height [3] (cm)** | **Ratio of throne height to leg length (%)** | **Distance between pedals (cm)** | **Pedal beater orientation (°)** | **Pedal board height (1-5)** | **Pedal spring tension (1-10)** |
| **1 – A** | 33 | 169 | 63 | 86 | 10.5 | 51.5 | 60 | 60 | 40 | 4 | 10 |
| **2 – A** | 25 | 185 | 70 | 97.5 | 11.7 | 54 | 55 | 42 | 65 | 3 | 10 |
| **3 – B** | 34 | 179 | 68 | 91 | 11.3 | 42.5 | 47 | 40 | 45 | 5 | 6 |
| **4 – B** | 32 | 173 | 75 | 90 | 12.5 | 50 | 56 | 63 | 50 | 3 | 8 |
| **5 – B** | 40 | 188 | 105 | 97 | 17.5 | 55 | 57 | 49 | 65 | 3 | 8 |
| **6 – B** | 28 | 182 | 85 | 97 | 14.2 | 60.5 | 62 | 42 | 45 | 3 | 8 |
| **7 – B** | 23 | 178 | 87 | 93.5 | 14.5 | 51 | 55 | 55 | 40 | 5 | 8 |
| **8 – A** | 23 | 188 | 70 | 96.5 | 11.7 | 58.5 | 61 | 52 | 35 | 2 | 3 |
| **9 – B** | 43 | 170 | 63 | 87.5 | 10.5 | 58.5 | 67 | 55 | 45 | 3 | 5 |
| **10 – B** | 36 | 166 | 57 | 84 | 9.5 | 45 | 54 | 57 | 40 | 3 | 7 |
| **11 – A** | 31 | 170 | 65 | 86.5 | 10.8 | 53.5 | 62 | 54 | 50 | 3 | 5 |
| **12 – B** | 44 | 178 | 95 | 90 | 15.8 | 54.5 | 61 | 55 | 45 | 2 | 8 |
| **13 – B** | 40 | 189 | 70 | 98.5 | 11.7 | 49.5 | 50 | 39 | 45 | 3 | 5 |
| **14 – B** | 39 | 180 | 80 | 94 | 13.3 | 56 | 60 | 40 | 30 | 1 | 7 |
| **15 – B** | 47 | 191 | 96 | 103 | 16.0 | 60 | 58 | 61 | 30 | 3 | 3 |
| **16 – A** | 35 | 170 | 70 | 87.5 | 11.7 | 56 | 64 | 51 | 35 | 4 | 8 |
| **17 – B** | 25 | 178 | 65 | 89.5 | 10.8 | 59 | 66 | 45 | 20 | 3 | 6 |
| **18 – A** | 42 | 186 | 79 | 98.5 | 13.2 | 59 | 60 | 42 | 30 | 3 | 7 |
| **Mean (SD)** | **34 (8)** | **179 (8)** | **76 (13)** | **92.6 (5.4)** | **12.6 (2.2)** | **54.1 (5.1)** | **59 (5)** | **50 (7)** | **42 (12)** | **3 (1)** | **7 (2)** |
| **Range** | **23-47** | **166-191** | **57-105** | **84-103** | **9.5-17.5** | **42.5-60.5** | **47-67** | **39-63** | **20-65** | **1-5** | **3-10** |

**Table S2.** Participating drummers' playing characteristics. Drummers' levels were categorized as follows: 'Regional' indicates performing frequently in shows, without participating in tours; 'National' indicates performing frequently in shows and participating to short national tours (< 15 consecutive shows); and 'International' indicates performing frequently in shows and participating in long international tours (> 15 consecutive shows). Drummers were grouped according to their maximal frequency (i.e., A: > 7.5 Hz; B: ≤ 7.5Hz).

| Drummer – group | Drumming Level [4] | Comfort frequency (Hz) | Maximal frequency (Hz) |
|---|---|---|---|
| **1 – A** | National | 6.33 | 7.7 |
| **2 – A** | Regional | 7.00 | 8.3 |
| **3 – B** | International | 6.00 | 7.3 |
| **4 – B** | National | 6.33 | 7.5 |
| **5 – B** | National | 6.00 | 7.0 |
| **6 – B** | Regional | 6.00 | 6.8 |
| **7 – B** | Regional | 6.00 | 6.6 |
| **8 – A** | Regional | 6.67 | 8.1 |
| **9 – B** | Regional | 5.67 | 7.0 |
| **10 – B** | Regional | 6.00 | 7.4 |
| **11 – A** | National | 6.00 | 7.9 |
| **12 – B** | International | 6.00 | 7.3 |
| **13 – B** | International | 6.00 | 7.2 |
| **14 – B** | National | 6.33 | 7.2 |
| **15 – B** | National | 6.33 | 6.9 |
| **16 – A** | International | 6.00 | 7.6 |
| **17 – B** | Regional | 6.33 | 6.7 |
| **18 – A** | National | 6.33 | 8.7 |
| **Mean (SD)** | | **6.18 (0.31)** | **7.0 (1.8)** |
| **Range** | | **5.67-7** | **6.6-8.7** |

**Table S3.** Drummers' individual results. Mean (SD) values for the ankle plantar flexion / dorsiflexion (FE) amplitude (°) and mean value (°), for the durations between onset and offset of SOL and of tibialis anterior (TA), and for the durations between offset of SOL or TA muscle and onset of TA or SOL muscle (ms), averaged for both sides, for the trial at 6.7 Hz and for the trial at the maximal frequency of each drummer. Durations for which mean – SD < 0 are displayed in bold and highlighted in red. Drummers were grouped according to their maximal frequency (i.e., A: > 7.5 Hz; B: ≤ 7.5Hz).

| | Ankle FE amplitude (°) | | Ankle FE mean value (°) | | Onset SOL- Offset SOL (ms) | | Onset TA – Offset TA (ms) | | Offset SOL - Onset TA (ms) | | Offset TA - Onset SOL (ms) | |
|---|---|---|---|---|---|---|---|---|---|---|---|---|
| **Drummer – group** | **6.7 Hz** | **Maximal frequency** | **6.7 Hz** | **Maximal frequency** | **6.7 Hz** | **Maximal frequency** | **6.7 Hz** | **Maximal frequency** | **6.7 Hz** | **Maximal frequency** | **6.7 Hz** | **Maximal frequency** |
| **1 – A** | 11.2 (0.7) | 8.0 (1.9) | 6.5 (1.2) | 8.4 (3.9) | 61.8 (6.1) | 66.3 (13.3) | 29.9 (12.7) | 58.9 (4.9) | 34.9 (8.7) | **2.5 (14.4)** | 14.9 (13.4) | **-4.3 (11.3)** |
| **2 – A** | 8.0 (0.5) | 4.4 (0.5) | 11.9 (1.8) | 4.4 (0.9) | 43.6 (14.6) | 50.0 (8.5) | NA | 40.5 (11.4) | NA | 24.8 (6.1) | NA | 13.7 (11.0) |
| **3 – B** | 11.8 (1.3) | 9.8 (0.9) | 1.8 (3.1) | -1.8 (2.2) | 58.6 (6.9) | 53.3 (5.9) | 53.5 (6.4) | 41.6 (8.8) | 18.7 (5.3) | 17.4 (7.1) | 17.7 (7.3) | 25.1 (8.3) |
| **4 – B** | 10.9 (1.8) | 8.1 (1.3) | -3.2 (2.6) | -6.8 (2.2) | 77.4 (8.3) | 74.8 (14.7) | 52.8 (10.0) | 36.4 (13.7) | 15.9 (13.5) | **11.9 (16.3)** | **5.2 (15.3)** | **5.8 (12.5)** |
| **5 – B** | 12.8 (0.8) | 8.6 (0.5) | -1.7 (0.7) | -2.0 (0.3) | 55.6 (7.1) | 51.4 (6.2) | 45.8 (5.0) | 42.5 (7.2) | 29.3 (7.3) | 34.5 (5.7) | 22.0 (6.6) | 16.9 (8.5) |
| **6 – B** | 9.1 (1.1) | 8.9 (0.5) | 0.8 (3.8) | -3.9 (2.9) | 50.3 (5.8) | 57.0 (8.7) | 33.8 (7.9) | 28.8 (8.5) | 36.1 (9.3) | 42.0 (12.6) | 30.8 (7.6) | 21.2 (8.0) |
| **7 – B** | 10.7 (2.1) | 10.7 (2.1) | -3.6 (5.4) | -3.6 (5.4) | 49.5 (7.9) | 49.5 (7.9) | 49.3 (4.4) | 49.3 (4.4) | 14.9 (10.0) | 14.9 (10.0) | 37.5 (10.1) | 37.5 (10.1) |
| **8 – A** | 6.5 (0.7) | 6.7 (1.4) | 1.1 (2.2) | 0.1 (2.6) | 37.0 (5.2) | 36.4 (7.0) | 37.5 (7.1) | 36.8 (11.1) | 32.6 (8.0) | 25.2 (10.2) | 28.4 (6.4) | 25.0 (6.1) |
| **9 – B** | 5.6 (1.7) | 5.4 (1.0) | 5.1 (0.7) | 3.7 (0.6) | 51.9 (8.6) | 53.7 (8.7) | 47.3 (6.0) | 48.4 (6.0) | 36.5 (7.4) | 32.2 (6.7) | 14.4 (8.0) | 10.6 (9.9) |
| **10 – B** | 4.9 (1.2) | 5.6 (2.0) | -2.4 (4.8) | -4.3 (5.1) | 56.9 (11.6) | 58.6 (7.6) | 60.2 (11.3) | 55.9 (16.2) | 29.9 (8.2) | 24.1 (13.5) | **3.6 (8.8)** | **-0.9 (10.3)** |
| **11 – A** | 7.0 (1.0) | 7.4 (2.2) | -1.8 (1.1) | -6.5 (1.1) | 51.1 (7.6) | 43.4 (10.1) | 43.0 (7.6) | 41.1 (6.6) | 28.9 (8.1) | 19.4 (6.6) | 27.0 (9.0) | 22.4 (5.7) |
| **12 – B** | 6.2 (2.4) | 4.7 (1.6) | -4.2 (2.0) | -7.5 (2.2) | 49.3 (7.5) | 41.1 (6.2) | 52.6 (5.8) | 48.8 (6.2) | 17.9 (5.4) | 19.4 (6.3) | 32.9 (5.2) | 27.9 (4.9) |
| **13 – B** | 7.4 (0.7) | 6.5 (0.5) | 23.5 (0.3) | 22.1 (0.4) | 47.4 (11.9) | 42.8 (10.1) | 93.4 (5.3) | 72.3 (6.9) | **-2.9 (13.1)** | **-2.4 (10.9)** | 11.7 (4.6) | 25.1 (8.3) |
| **14 – B** | 6.3 (0.8) | 5.9 (1.0) | -4.8 (0.5) | -7.1 (0.5) | 47.4 (4.1) | 49.5 (8.0) | 52.0 (6.5) | 52.8 (6.6) | 27.2 (6.1) | 26.6 (9.2) | 19.8 (6.6) | 12.0 (7.9) |
| **15 – B** | 2.5 (0.8) | 2.5 (1.1) | -10.7 (7.3) | -11.8 (10.2) | 52.2 (6.0) | 51.1 (7.8) | 48.1 (10.2) | 43.9 (11.3) | 28.8 (12.6) | 28.2 (13.2) | 22.0 (6.9) | 21.9 (9.4) |
| **16 – A** | 11.0 (2.3) | 9.1 (2.4) | -6.8 (0.9) | -5.4 (1.6) | 48.4 (5.9) | 48.0 (7.2) | 30.1 (10.7) | 33.6 (12.7) | 42.7 (10.0) | 34.7 (13.1) | 23.0 (10.4) | 22.9 (4.8) |
| **17 – B** | 6.4 (1.0) | 6.4 (1.0) | 8.1 (2.2) | 8.1 (2.2) | 40.3 (3.6) | 40.3 (3.6) | 50.1 (6.5) | 50.1 (6.5) | 24.4 (5.0) | 24.4 (5.0) | 32.7 (6.9) | 32.7 (6.9) |
| **18 – A** | 10.8 (4.6) | 6.2 (0.8) | -2.4 (1.1) | -7.1 (1.0) | 57.9 (12.1) | 53.9 (10.7) | 30.9 (11.3) | 41.1 (7.5) | **22.8 (23.7)** | **-2.6 (7.0)** | 37.8 (4.5) | NA |
| **Mean (SD)** | **7.8 (3.4)** | **6.9 (2.1)** | **1.0 (7.9)** | **-1.2 (8.0)** | **52.0 (8.9)** | **51.2 (9.3)** | **47.7 (15.0)** | **45.7 (10.2)** | **25.8 (10.8)** | **20.8 (12.4)** | **22.4 (10.4)** | **18.6 (11.2)** |
| **Range** | **2.5-12.8** | **2.5-10.7** | **-10.7-23.5** | **-11.8-22.1** | **37.0-77.4** | **36.4-74.8** | **29.9-93.4** | **28.8-72.3** | **-2.9-42.7** | **-2.6-42.0** | **3.6-37.8** | **-4.3-37.5** |

**Table S4.** Drummers' individual results. Mean (SD) values for the duration between onset of ankle dorsiflexion and onset of SOL, for the duration between onset of SOL and ankle plantar flexion, for the duration between ankle plantar flexion and onset of tibialis anterior (TA), and for the duration between onset of TA and ankle dorsiflexion, averaged for both sides, for the trial at 6.7 Hz and for the trial at the maximal frequency of each drummer. Drummers were grouped according to their maximal frequency (i.e., A: > 7.5 Hz; B: ≤ 7.5Hz).

| | Ankle dorsiflexion – Onset SOL (ms) | | Onset SOL- Ankle plantar flexion (ms) | | Ankle plantar flexion – Onset TA (ms) | | Onset TA – Ankle dorsiflexion (ms) | |
|---|---|---|---|---|---|---|---|---|
| **Drummer – group** | **6.7 Hz** | **Maximal frequency** | **6.7 Hz** | **Maximal frequency** | **6.7 Hz** | **Maximal frequency** | **6.7 Hz** | **Maximal frequency** |
| **1 – A** | 37.6 (2.0) | 30.3 (12.1) | 39.4 (3.1) | 37.6 (12.5) | 58.4 (6.9) | 28.4 (7.4) | 11.9 (6.1) | 34.1 (7.0) |
| **2 – A** | 36.3 (2.3) | 31.5 (5.9) | 38.2 (2.5) | 34.8 (6.0) | NA | 32.5 (5.3) | NA | 21.3 (6.6) |
| **3 – B** | 47.6 (4.3) | 47.6 (6.5) | 32.8 (3.4) | 26.2 (3.3) | 44.6 (4.0) | 44.2 (5.1) | 25.1 (4.0) | 18.5 (3.9) |
| **4 – B** | 44.8 (3.5) | 39.6 (8.2) | 38.2 (4.3) | 30.3 (11.4) | 50.5 (8.7) | 47.4 (6.6) | 11.6 (9.6) | 11.5 (5.0) |
| **5 – B** | 53.6 (4.5) | 57.6 (3.2) | 22.8 (3.1) | 22.1 (3.1) | 62.1 (5.9) | 63.2 (3.0) | 13.1 (4.1) | 1.8 (2.8) |
| **6 – B** | 42.1 (2.8) | 37.8 (6.3) | 39.4 (4.2) | 42.3 (7.1) | 47.1 (11.5) | 56.5 (12.4) | 23.5 (8.3) | 11.7 (10.2) |
| **7 – B** | 55.1 (12.5) | 55.1 (12.5) | 38.4 (5.0) | 38.4 (5.0) | 26.0 (10.5) | 26.0 (10.5) | 31.6 (8.0) | 31.6 (8.0) |
| **8 – A** | 49.1 (5.4) | 43.8 (6.7) | 27.1 (5.1) | 26.0 (5.5) | 42.5 (7.6) | 35.9 (12.6) | 17.6 (7.0) | 18.7 (14.7) |
| **9 – B** | 29.9 (7.0) | 24.1 (8.0) | 50.1 (6.6) | 51.0 (6.4) | 37.4 (6.1) | 34.9 (5.6) | 31.4 (7.2) | 34.8 (7.8) |
| **10 – B** | 27.0 (14.7) | 18.3 (8.4) | 39.7 (7.5) | 44.7 (6.3) | 47.0 (11.3) | 38.1 (10.9) | 36.4 (12.5) | 36.4 (12.1) |
| **11 – A** | 58.3 (5.2) | 48.3 (7.2) | 25.8 (5.3) | 21.3 (4.7) | 54.4 (6.5) | 41.4 (8.2) | 11.4 (4.9) | 15.8 (6.6) |
| **12 – B** | 36.8 (15.2) | 42.4 (8.3) | 41.0 (6.6) | 34.4 (6.4) | 26.3 (11.0) | 25.8 (8.6) | 47.5 (8.8) | 34.2 (7.8) |
| **13 – B** | 80.9 (5.2) | 71.8 (5.9) | 28.8 (5.1) | 23.7 (6.2) | 16.4 (5.4) | 16.3 (4.4) | 23.9 (3.9) | 24.5 (3.2) |
| **14 – B** | 42.8 (5.4) | 39.1 (6.1) | 39.3 (3.7) | 36.6 (5.7) | 35.3 (6.5) | 36.4 (8.8) | 28.9 (5.8) | 25.4 (6.4) |
| **15 – B** | 34.0 (11.4) | 34.9 (12.7) | 41.3 (6.8) | 31.1 (12.9) | 39.8 (13.3) | 48.1 (15.2) | 37.7 (9.7) | 31.1 (14.8) |
| **16 – A** | 37.6 (4.2) | 38.4 (7.8) | 43.9 (5.6) | 37.6 (5.8) | 45.4 (7.8) | 45.7 (15.7) | 21.3 (7.7) | 14.9 (11.9) |
| **17 – B** | 49.9 (8.3) | 49.9 (8.3) | 41.2 (6.7) | 41.2 (6.7) | 22.9 (6.7) | 22.9 (6.7) | 33.9 (8.8) | 33.9 (8.8) |
| **18 – A** | 44.7 (6.3) | 33.0 (8.1) | 35.8 (7.8) | 29.4 (6.4) | 44.8 (17.6) | 26.1 (4.5) | 25.3 (10.5) | 26.9 (4.5) |
| **Mean (SD)** | **44.9 (12.4)** | **41.3 (12.7)** | **36.8 (6.9)** | **33.8 (8.2)** | **41.2 (12.6)** | **37.2 (12.2)** | **25.4 (10.4)** | **23.7 (10.0)** |
| **Range** | **27.0-80.9** | **18.3-71.8** | **22.8-50.1** | **21.3-51.0** | **16.4-62.1** | **16.3-63.2** | **11.4-47.5** | **1.8-36.4** |

**Table S5.** Drummers' muscle activation individual results. Mean (SD) values for the 90th percentile of muscle activation in percent of the Maximal Voluntary Contraction (MVC), averaged for both sides, for the trial at 6.7 Hz and for the trial at the maximal frequency of each drummer. Activations for which mean + SD > 30% MVC are in bold and highlighted in red. ES: Erector Spinae; GM: Gluteus Medius; RA: Rectus Abdominis; RF: Rectus Femoris; TFL: Tensor Fascia Latae. Drummers were grouped according to their maximal frequency (i.e., A: > 7.5 Hz; B: ≤ 7.5Hz).

| | 90th percentile ES (% MVC) | | 90th percentile GM (% MVC) | | 90th percentile RA (% MVC) | | 90th percentile RF (% MVC) | | 90th percentile TFL (% MVC) | |
|---|---|---|---|---|---|---|---|---|---|---|
| **Drummer – group** | **6.7 Hz** | **Maximal Frequency** | **6.7 Hz** | **Maximal Frequency** | **6.7 Hz** | **Maximal Frequency** | **6.7 Hz** | **Maximal Frequency** | **6.7 Hz** | **Maximal Frequency** |
| **1 – A** | 9.1 (4.5) | **44.9 (17.8)** | 8.5 (5.7) | **31.9 (12.5)** | 3.2 (2.9) | 8.2 (5.7) | 11.3 (6.4) | **54.0 (19.7)** | 11.6 (9.6) | **28.9 (14.5)** |
| **2 – A** | 23.2 (3.8) | **24.1 (9.3)** | 8.8 (2.9) | **44.3 (11.9)** | NA | NA | 5.1 (0.3) | 15.1 (4.9) | 6.3 (3.0) | 12.1 (8.0) |
| **3 – B** | 2.0 (1.0) | 1.5 (0.5) | 10.9 (2.2) | 19.6 (3.9) | 1.1 (0.5) | 1.2 (0.5) | 10.8 (2.1) | 15.2 (2.8) | 9.7 (2.0) | 16.3 (4.5) |
| **4 – B** | 12.1 (5.6) | 16.9 (7.4) | 14.7 (5.8) | **30.4 (17.4)** | 5.3 (3.1) | 5.8 (3.2) | 6.4 (4.2) | 11.3 (9.8) | **24.1 (5.3)** | **34.1 (8.4)** |
| **5 – B** | 14.0 (5.3) | 18.1 (7.7) | 15.7 (2.7) | 18.5 (6.5) | 13.6 (4.2) | 12.9 (3.5) | 7.1 (0.8) | 9.5 (3.6) | 11.4 (4.7) | **34.2 (15.6)** |
| **6 – B** | 8.0 (5.0) | 6.7 (3.6) | 4.9 (1.8) | 5.0 (3.1) | 11.9 (10.4) | 16.0 (10.7) | **20.9 (16.5)** | **18.5 (13.4)** | 11.9 (1.8) | 18.0 (3.9) |
| **7 – B** | **39.8 (9.0)** | **39.8 (9.0)** | **38.2 (22.2)** | **38.2 (22.2)** | 15.7 (6.1) | 15.7 (6.1) | **42.3 (18.8)** | **42.3 (18.8)** | **32.4 (10.1)** | **32.4 (10.1)** |
| **8 – A** | **27.5 (7.2)** | **35.7 (8.2)** | 1.9 (0.8) | 3.8 (3.9) | 1.9 (1.7) | 2.2 (1.5) | **36.0 (13.2)** | **69.3 (21.5)** | 5.1 (2.4) | 10.5 (4.1) |
| **9 – B** | 16.5 (7.8) | 15.5 (7.2) | 3.2 (1.9) | 9.7 (2.9) | 4.7 (3.8) | 7.3 (3.4) | 9.8 (4.3) | 16.9 (6.6) | 7.4 (5.3) | 16.1 (5.0) |
| **10 – B** | 18.7 (4.4) | 17.8 (4.8) | 8.3 (1.9) | 14.9 (6.0) | 1.7 (1.3) | 2.1 (1.4) | 3.6 (0.7) | 7.6 (1.8) | **29.1 (13.4)** | **44.5 (31.6)** |
| **11 – A** | 8.2 (4.9) | 10.9 (4.0) | 4.9 (3.9) | **33.7 (34.2)** | 1.9 (1.3) | 2.1 (1.1) | 9.6 (2.5) | **23.0 (7.3)** | 4.3 (3.8) | 10.5 (6.4) |
| **12 – B** | **21.9 (9.2)** | **30.3 (14.5)** | **31.3 (9.0)** | **44.6 (17.0)** | 11.7 (6.4) | **22.3 (9.6)** | **60.8 (37.6)** | **99.4 (32.7)** | **38.5 (23.8)** | **60.8 (36.2)** |
| **13 – B** | 18.5 (7.5) | **20.6 (10.3)** | **41.3 (23.4)** | **49.9 (31.6)** | 13.6 (6.1) | 11.9 (4.7) | **36.9 (7.1)** | **42.7 (11.1)** | **58.0 (13.2)** | **57.9 (14.7)** |
| **14 – B** | 7.5 (5.1) | 10.0 (7.6) | 3.9 (1.7) | 10.1 (3.1) | 2.2 (1.7) | 2.6 (1.7) | 11.5 (8.6) | 16.8 (8.3) | 2.4 (1.0) | 9.6 (6.6) |
| **15 – B** | 4.5 (2.6) | 5.2 (2.1) | 12.9 (3.7) | 16.7 (7.2) | 6.3 (3.0) | 9.6 (5.2) | 14.1 (11.1) | 15.2 (12.8) | 20.5 (6.0) | 22.6 (5.5) |
| **16 – A** | 5.8 (4.0) | 11.5 (9.8) | 10.6 (9.9) | **61.6 (53.4)** | 0.9 (0.8) | 0.9 (0.7) | 7.2 (1.6) | 19.0 (8.5) | 3.9 (3.8) | 11.5 (8.6) |
| **17 – B** | 6.5 (2.4) | 6.5 (2.4) | 5.4 (4.2) | 5.4 (4.2) | 2.0 (1.3) | 2.0 (1.3) | 12.6 (4.6) | 12.6 (4.6) | 6.6 (3.0) | 6.6 (3.0) |
| **18 – A** | 5.2 (4.1) | 17.5 (10.5) | 13.7 (15.8) | **36.7 (31.6)** | 1.1 (1.0) | 8.5 (9.4) | 16.0 (11.9) | **77.7 (26.7)** | 5.8 (3.2) | **27.7 (8.8)** |
| **Mean (SD)** | **13.8 (9.8)** | **18.5 (12.3)** | **13.3 (11.7)** | **26.4 (17.2)** | **5.8 (5.3)** | **7.7 (6.3)** | **17.9 (15.7)** | **31.5 (27.0)** | **16.1 (15.0)** | **25.2 (16.3)** |
| **Range** | **2.0-39.8** | **1.5-44.9** | **1.9-41.3** | **3.8-61.6** | **0.9-15.7** | **0.9-22.3** | **3.6-60.8** | **7.6-99.4** | **2.4-58.0** | **6.6-60.8** |
